# Exogenous lipid vesicles induce endocytosis-mediated cellular aggregation in a close unicellular relative of animals

**DOI:** 10.1101/2024.05.14.593945

**Authors:** Ria Q. Kidner, Eleanor B. Goldstone, Henry J. Rodefeld, Lorin P. Brokaw, Aria M. Gonzalez, Núria Ros-Rocher, Joseph P. Gerdt

**Author notes:** **Corresponding author:** Joseph P. Gerdt.

## Abstract

*Capsaspora owczarzaki* is a protozoan that may both reveal aspects of animal evolution and also curtail the spread of schistosomiasis, a neglected tropical disease. *Capsaspora* exhibits a chemically regulated aggregative behavior that resembles cellular aggregation in some animals. This behavior may have played a key role in the evolution of animal multicellularity. Additionally, this aggregative behavior may be important for *Capsaspora*’s ability to colonize the intermediate host of parasitic schistosomes and potentially prevent the spread of schistosomiasis. Both applications demand elucidation of the molecular mechanism of *Capsaspora* aggregation. Toward this goal, we first determined the necessary chemical properties of lipid cues that activate aggregation. We found that a wide range of abundant zwitterionic lipids induced aggregation, revealing that the aggregative behavior could be activated by diverse lipid-rich conditions. Furthermore, we demonstrated that aggregation in *Capsaspora* requires clathrin-mediated endocytosis, highlighting the potential significance of endocytosis-linked cellular signaling in recent animal ancestors. Finally, we found that aggregation was initiated by post-translational activation of cell-cell adhesion—not transcriptional regulation of cellular adhesion machinery. Our findings illuminate the chemical, molecular and cellular mechanisms that regulate *Capsaspora* aggregative behavior—with implications for the evolution of animal multicellularity and the transmission of parasites.

## INTRODUCTION

*Capsaspora owczarzaki* is a protozoan that may inform the origins of animal multicellularity and also curtail the spread of a neglected tropical disease. From the evolutionary perspective, *Capsaspora* is one of the closest living relatives of animals, and its genome shares many genes involved in animal cell-cell signaling and adhesion.^1, 2^ Therefore genomic comparisons between extant animals, *Capsaspora*, and other close unicellular relatives of animals reveal the ancestry and evolution of genes that are essential for animal multicellularity.^2^ Furthermore, *Capsaspora* was isolated from the snails that transmit the parasitic worms that cause schistosomiasis.^3, 4^ *Capsaspora* can kill these parasites in their intramolluscan life stage, potentially preventing human infections.^3, 5, 6^ Notably, *Capsaspora* exhibits a multicellular behavior that is likely important *both* for studying the origins of animal multicellularity and for studying the potential of *Capsaspora* to combat schistosomiasis. Namely, *Capsaspora* forms large multicellular aggregates in response to lipids from its host snail.^7, 8^

In the context of animal multicellularity, cellular aggregation may have been an important part of the life cycle of the first multicellular animal.^7, 9, 10^ Today, reversible cellular aggregation remains an important behavior for many animal cells (e.g., whole body regeneration in sponge cells,^11^ cellular ingression during gastrulation of mammalian epiblasts,^12^ migration of neuronal cells in developing nervous systems,^13^ and immune cell aggregation during inflammation^14^). Yet it is still unclear which aspects of the reversible aggregation behaviors in present-day animals are homologous to the morphologically similar behaviors in *Capsaspora*^2, 7^ or other unicellular animal relatives.^9, 15, 16^ Specifically, homologous genes driving similar behaviors in *Capsaspora* may imply ancient ancestry of aspects of aggregative multicellularity in animals.^2^ Therefore, we must better understand the molecular mechanisms of *Capsaspora* aggregation induction in order to compare it to similar animal behaviors.

In the context of schistosomiasis transmission, *Capsaspora* aggregation may be important for colonization of host snails. Since *Capsaspora* likely must be inside the vector snails to prevent the maturation and spread of schistosomes, it is essential to understand how *Capsaspora* senses and adapts to its host environment. Since it aggregates within snail tissue (and in response to specific snail-derived lipids),^8^ we suspect this response is important for host colonization. However, we still poorly understand the regulatory mechanism by which *Capsaspora* senses these host factors.

For both of these reasons, we aim to elucidate the molecular and cellular mechanisms that regulate multicellular aggregation in *Capsaspora*. Despite significant progress in characterizing this phenomenon,^2, 7, 17^ the mechanism(s) underlying the initial induction and maintenance of aggregation in *Capsaspora* remains elusive.

Building upon previous work demonstrating that low-density lipoproteins (LDLs), snail-derived lipids, and pure dioleoylphosphatidylcholine (DOPC) induced *Capsaspora* aggregation,^7, 8^ we investigated which other pure lipids induced aggregation. We examined several natural lipids present in LDLs and snail serum, as well as non-native lipids, to discern the lipid properties necessary to induce aggregation. We found that many zwitterionic diacyl lipids robustly induced aggregation. This finding indicates that *Capsaspora* (and possibly its ancestors) can sense a wide range of universally abundant lipids that could serve as general cues for environments with nutritional prey.

Furthermore, we investigated the mechanism of action of these inducers within the *Capsaspora* cell. Given the established importance of receptor-mediated endocytosis in LDL uptake by human cells, we hypothesized a similar role in *Capsaspora* aggregation. Indeed, our findings demonstrated that endocytosis is necessary for aggregation, as chemical inhibition of various early steps in the endocytic pathway prevents this process. In addition, we showed that initiation of aggregation occurs within seconds of lipid addition and does not rely on translation of new proteins. Nevertheless, subsequent transcriptionally regulated processes could likely promote further development of the aggregates.^2^

In summary, our study sheds light on the lipid-mediated induction of cellular aggregation in *Capsaspora* and highlights the crucial role of endocytosis and post-translational regulation in this process. These insights not only advance our understanding of unicellular-to-multicellular transitions but also provide insight into a behavior that may be important for a protective symbiosis.

## RESULTS

### Phosphatidylcholine-mediated induction of *Capsaspora* multicellularity correlates with lipid fluidity

In previous work, we found that fetal bovine serum (FBS), low density lipoproteins (LDLs), snail serum, and pure dioleoyl phosphatidylcholine (DOPC) lipids induced robust cellular aggregation in *Capsaspora*.^7, 8^ However, we suspected that other natural lipids beyond DOPC would also induce aggregation for a few reasons: A) many lipids are structurally similar to DOPC, B) the concentration of DOPC present in snail serum was insufficient to fully account for the activity of aggregation induction by crude snail serum,^8^ and C) we found that HDLs (which contain only minor amounts of DOPC)^18^ also robustly induced aggregates (**Fig. 1A–B**). To determine which other phospholipids induced *Capsaspora* aggregation, we began a structure-activity relationship (SAR) study for common phospholipids. We first determined the importance of macromolecular lipid assembly for activity, followed by the requirements in the polar headgroup, and finally the significance of the lipophilic tail composition.

**Figure 1:**
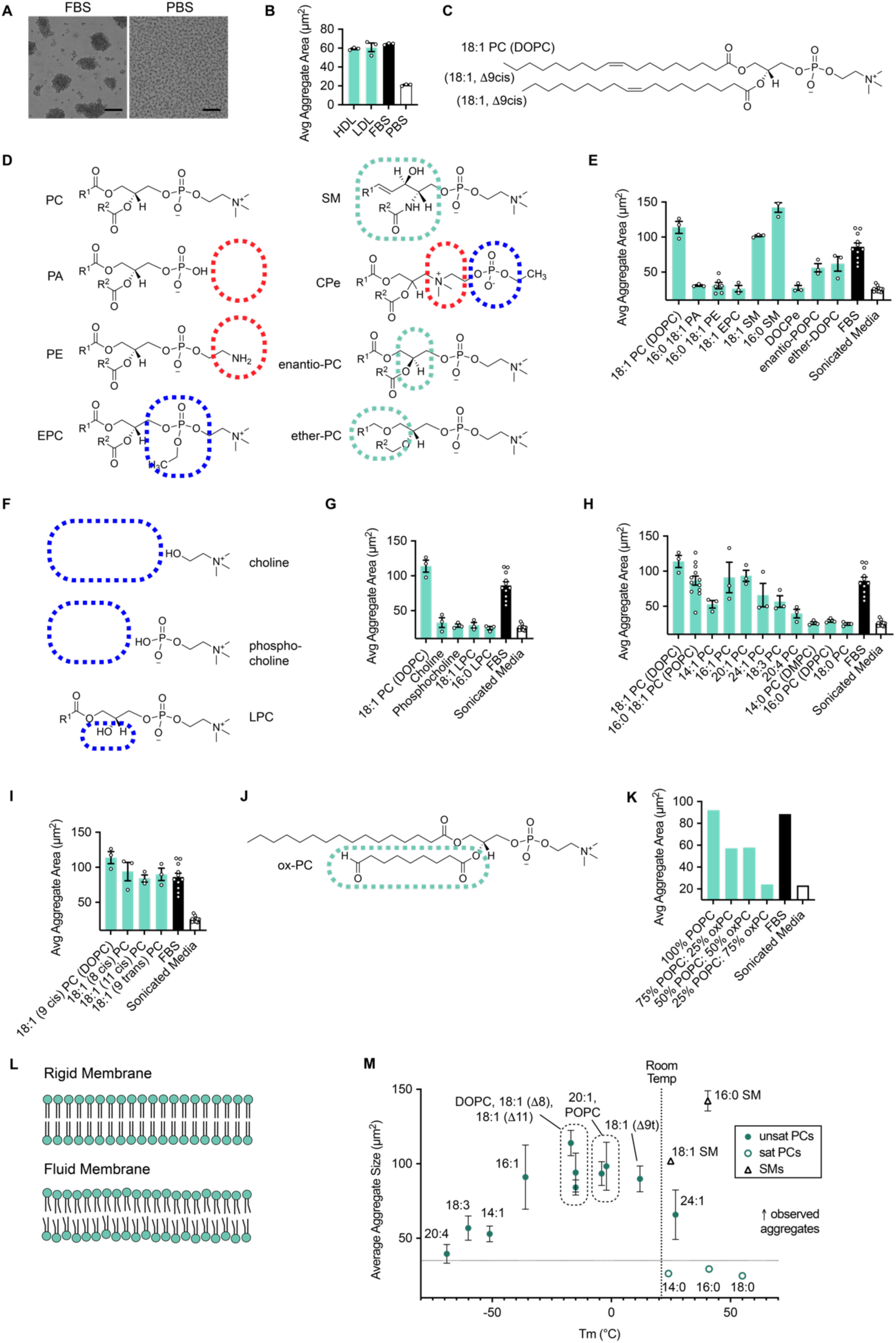
Structure activity relationship of lipids that induce aggregation in *Capsaspora*. (A) Representative images of *Capsaspora* cellular aggregates induced by 5% (v/v) FBS compared to single cells with 5% (v/v) PBS added. Scale bar is 50 µm. (B) Plot of average aggregate area induced by 80 µg/mL of high density lipoprotein (HDL) and low density lipoprotein (LDL), compared to controls 5% (v/v) FBS and 5% (v/v) PBS. HDL and LDL are active. (C) Chemical structure of DOPC, the previously reported active phospholipid.^8^ (D) Head group structures for tested lipids. (E) Plot of aggregation induction activity of lipids with varying head groups. Variations in lipid monomer head groups show that aggregation-induction activity is unique to phosphatidylcholine (PC) and sphingomyelin (SM) head groups. (F) Structures of choline molecules lacking lipophilic tails. (G) Plot of aggregation induction activity of lipids with different numbers of tails, ranging from no tails to two tails. Phosphatidylcholine lipid monomers with two tails are active while lyso-PCs and phosphocholine/choline alone are inactive. (H) Plots of aggregation resulting from PCs with different tail lengths and degrees of unsaturation. PCs with at least one degree of unsaturation were active while no saturated PCs induced aggregation. (I) Plots of aggregation resulting from induction with PCs with different positions and stereochemistry of unsaturation. The geometry and location of the lipid tail double bond does not influence aggregate-inducing activity. (J) The structure of oxPC, an oxidative lipid product identified in vesicle preparations. (K) Aggregation induced by mixtures of POPC with oxPC. Addition of more oxPC did not cause more activity. (L) Cartoon of lipid membrane fluidity due to increased points of unsaturation. (M) Aggregation resulting from previously tested PCs and SMs plotted as a function of lipid gel phase transition temperature.^20^ Activity of PC vesicles is generally associated with a gel phase transition temperature close to or below room temperature. In all aggregation plots, error bars represent standard error of the mean. Individual values are shown as open circles. For each inducer, multiple concentrations were tested (**Fig. S2**) and the most active concentration is shown here. Single images were analyzed for panel (K).

First, we found that lipids must be formed into vesicles prior to inducing aggregation activity. Lipids that were simply mixed with buffer and delivered to cells did not induce activity, but even brief sonication formed vesicles that induced robust aggregation (**Fig. S1A**). Next, we tested how the chemical composition of the lipids within these vesicles influenced aggregation (**Fig. 1C–M**, **S2**). To ensure that vesicles formed correctly, each lipid in the SAR study was characterized by TEM (**Fig. S1B–U**). Since DOPC was previously reported to be active,^8^ we used the chemical structure of DOPC as our reference (**Fig. 1C**). Since phospholipids are classically categorized by the composition of their polar headgroups and lipophilic tails, we systematically varied these components and tested for aggregation induction. We first varied the structures of headgroups (**Fig. 1D–E**). Lipids with phosphatidylcholine (PC) headgroups were most active compared to lipids with just a phosphate (i.e., phosphatidic acid, PA) or phosphatidyl ethanolamine (PE). Therefore, the permanent positive charge afforded by the choline appears to be important for aggregation activity. The negative charge of the phosphodiester also was essential, evidenced by the failure of the matching ethylphosphocholine (EPC) lipid to activate aggregation. Therefore, a zwitterion is necessary for aggregation activity in *Capsaspora*. The importance of this zwitterion is further supported by the aggregation-inducing activity of sphingomyelin (SM) lipids. Intriguingly, the order of the positive and negative charges on the zwitterionic head group is essential. As evidenced by the inactivity of DOCPe, the lipid positive charge must be exterior to the negative charge. As suggested by the activity of the SM lipids, the glycerol ester moieties in PCs can be modified without losing activity. For example, the chirality of the glycerophophocholine headgroup did not control activity—the enantiomeric PC still induced aggregation. Furthermore, a diether PC was active. Therefore, to induce aggregation, the lipids must be assembled into soluble nanoscale particles, and they must possess a zwitterionic headgroup with the positive charge facing outward.

Next, we determined which features of the phospholipid tails were necessary for activity. First, we found that two lipophilic tails were required for activity. The simple addition of choline or phosphocholine failed to induce aggregation (**Fig. 1F–;G**). Furthermore, a single lipid tail was insufficient for activity. Each lysophosphatidylcholine (LPC) tested was inactive (**Fig. 1G**). Therefore, only diacyl zwitterionic lipids induce aggregation. At least one of the two tails also needed at least one degree of unsaturation to be active (**Fig. 1H**). However, there was no strict requirement for the geometry or location of the double bond. Lipids with double bonds further or closer to the headgroup retained activity, and lipids with either cis or trans olefins were equally active (**Fig. 1I**). We considered two possible explanations to the necessity of a double bond. First, the alkene may be oxidized in handling,^19^ and the oxidized product is active. Alternatively, the increased fluidity afforded by a double bond is essential for activity. Moving forward, we tested the validity of each of these hypotheses.

It has been shown that PCs with double bonds can undergo oxidative cleavage (**Fig. 1J**). ^19^ We used LCMS to determine if oxidative cleavage products were present in our lipid vesicles, and indeed, we discovered an oxidized PC (hereafter oxPC) was present at levels ~1000x higher in the prepared vesicles than in original storage vials (**Fig. S3**). To test if the activity was due to the presence of these oxidized products, we added extra oxPC to prepared POPC vesicles. We found that additional oxPC did not increase the potency of the lipid vesicles. In fact, these vesicles were less active (**Fig. 1K**). This result suggests that lipid oxidative cleavage is not the primary reason that one degree of unsaturation is required in the inducer.

Next, we considered the importance of lipid fluidity for inducing aggregation. Classically, saturated lipids form more rigid lipid bilayers while unsaturated lipids form more fluid bilayers (**Fig. 1L**). Since gel phase transition temperature of phospholipids correlates with fluidity,^20^ we plotted all tested lipids in order of reported gel phase transition temperature^20^ to determine if physical fluidity of vesicles was correlated with activity (**Fig. 1M**). Indeed, all active lipids with a phosphatidylcholine headgroup and two tails were active as long as the gel phase transition temperature laid below room temperature (**Fig. 1M**). This room temperature cutoff is not universal across all lipids, though, since an active sphingomyelin [SM(16:0)] has a Tm ~40 °C.

Overall, the necessary characteristics for an aggregation-inducing lipid are A) delivery as vesicles, B) a zwitterionic headgroup with the positive charge exposed to the surface, C) two tails with at least one degree of unsaturation, and D) for PCs, a gel phase transition temperature below room temperature.

### Induction of *Capsaspora* multicellular aggregates is dependent on clathrin-mediated endocytosis

We previously observed that *Capsaspora* depleted LDL from the media, and it appeared to accumulate LDL within its cells.^7^ Since low density lipoprotein (LDL) is taken up by human cells through receptor mediated endocytosis (**Fig. 2A**),^21^ we hypothesized that this pathway was involved in *Capsaspora* lipoprotein uptake as well. We first monitored the uptake of a fluorescently-labelled LDL and found that it accumulated inside cells in small puncta (**Fig. 2B**). We then doped a fluorescently-labelled PC into pure DOPC vesicles and saw similar intracellular accumulation (**Fig. 2C**). LDL with the pH-dependent dye “pHrodo red” fluoresced upon cell uptake, suggesting that the LDL accumulates inside cells in vesicles with low pH^22^ (*e.g.*, endosomes and lysosomes) (**Fig. 2D**). Therefore, LDL and PC particles are endocytosed by *Capsaspora*.

**Figure 2:**
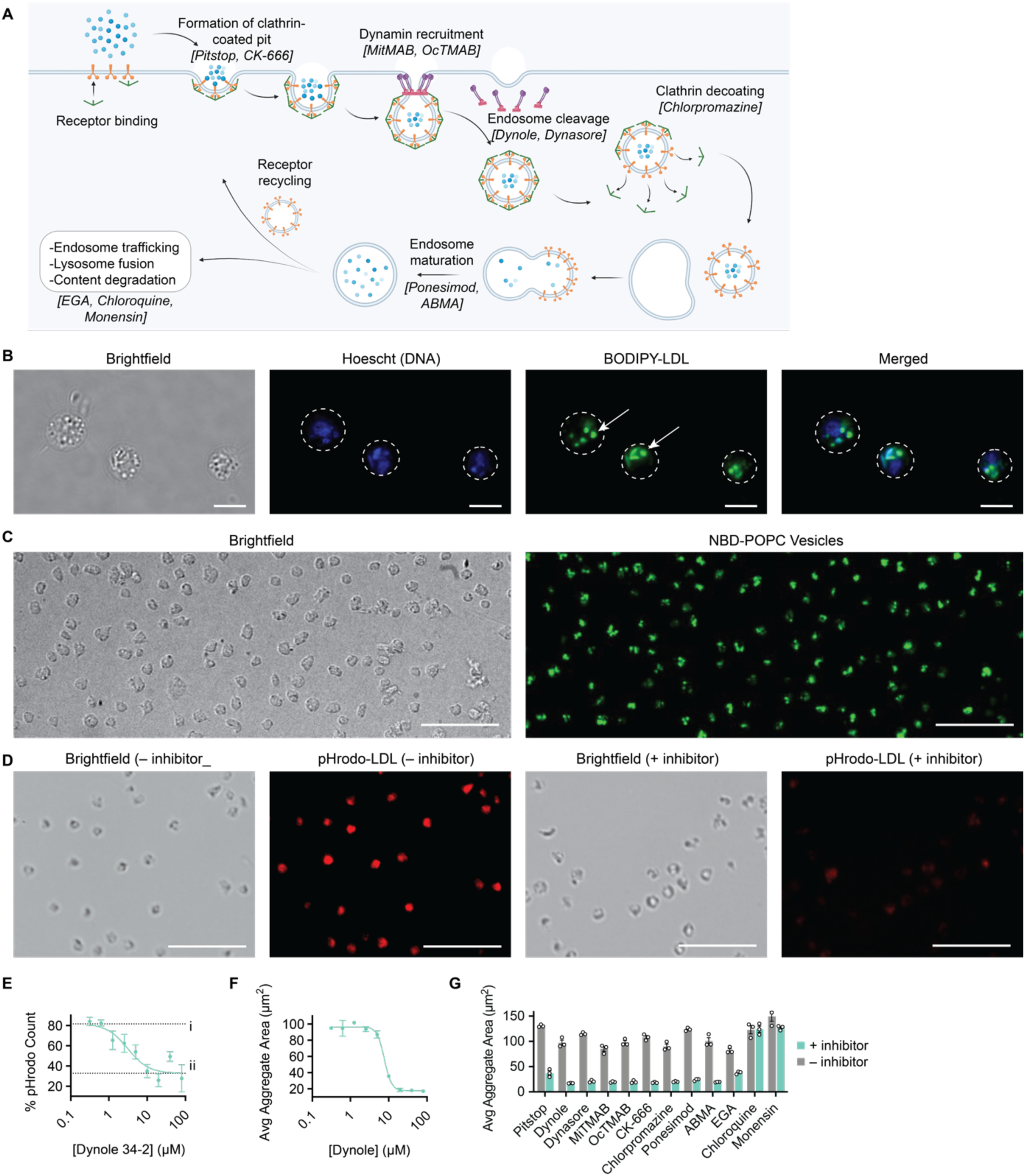
Aggregation is dependent on receptor mediated endocytosis of LDL and POPC. (A) Stages of clathrin-mediated endocytosis, labeled in brackets with the pharmacological inhibitors used to target each stage. (B) Cellular uptake of BODIPY labelled LDL shows accumulation of fluorescent puncta in adherent cells (puncta highlighted with white arrows). Scale bars are 5 µm. (C) Uptake of POPC doped at 5% (w/w) with NBD-PC shows similar results to LDL and accumulation of puncta inside adherent cells. Scale bars are 50 µm. (D) Uptake of pHrodo-LDL shows accumulation of red fluorescent puncta in acidified endosomes/lysosomes, which is prevented by incubation with an endocytosis inhibitor (here, Dynole 34-2). Scale bars are 50 µm. (E) Percent of adherent cells with pHrodo red fluorescence after inhibition of LDL uptake with the dynamin inhibitor Dynole 34-2. The inhibitor blocked fluorescent pHrodo LDL accumulation. (F) Average aggregate area of cells treated with a dilution series of the inhibitor Dynole 34-2. Aggregation was induced by FBS 30 minutes after inhibitor treatment. The dynamin inhibitor Dynole 34-2 inhibited aggregation with a potency that correlated with its inhibition of LDL uptake. (G) Average aggregate area upon treatment with various chemical inhibitors along the receptor mediated endocytosis pathway (See **Fig. S4** for complete dose-response curves). Aggregates were induced with FBS 30 minutes after treatment with each inhibitor. Grey bars indicate DMSO controls tested concurrently with the matching inhibitor treatment. Consistent aggregation inhibition along the pathway shows that the initial stages of endocytosis are necessary for aggregation. In all plots, error bars represent standard error of the mean (n=3). Panel A was created with BioRender.com.

To determine if endocytosis of the lipid particles was necessary for aggregation, we used a chemical genetics approach to block different steps of the endocytosis pathway. First, we found that the dynamin inhibitor Dynole 34-2, inhibited *both* uptake of fluorescently labelled particles (**Fig. 2D–E**) *as well as* aggregate formation in a dose-dependent manner (**Fig. 2F**) with similar IC50 values ~5 µM. With this initial result, we then systematically tested several steps of the endocytosis pathway with other chemical inhibitors (**Fig. 2A, G**, **Fig. S4**). Inhibition of clathrin recruitment to the cell surface (using Pitstop 2) prevented aggregation in *Capsaspora*. Actin polymerization and branching is necessary for endosome budding into the clathrin pit at the cell surface. Inhibition of actin branching (using CK-666) also prevented cellular aggregation.^23^ Inhibition of dynamin binding (using either MiTMAB or OcTMAB) as well as dynamin activation (using either Dynasore or Dynole 34-2) prevented aggregation at concentrations comparable to those used in animal cell culture studies.^24–26^ Inhibiting clathrin de-coating of the early endosome (using chlorpromazine) also prevented aggregate formation.^27^ Inhibiting endosome maturation from the early endosome to the late endosome (using either Ponesimod or ABMA) also prevented formation of aggregates suggesting that inducers must reach the late endosome for proper formation of aggregates.^28, 29^ On the other hand, inhibiting vesicle trafficking to the endolysosome (using EGA)^30^ failed to completely prevent aggregation, even at very high concentrations. Additionally, inhibiting lysosome acidification (using either Chloroquine or Monensin)^31, 32^ failed to prevent aggregation. Taken together, these results strongly suggest that receptor mediated endocytosis is essential for cellular aggregation in *Capsaspora*. Although lipid particles accumulate in acidified lysosomes, it does not appear that the late stages of lysosome maturation are required for cellular aggregation. Only the earlier steps of the endocytic pathway appear to be required to activate *Capsaspora* aggregation.

### Initial cellular aggregation is activated post-translationally within seconds

In previous work, we observed aggregation initiate within three minutes of lipid addition, after which the aggregates continued to coalesce over the course of several hours before dissipating.^7^ With shorter time-series analyses, we now observed cells start to aggregate within seconds after addition of inducer (**Movie 1, Fig. 3A**). This immediate aggregation is also apparent after addition of fluorescent PC lipid particles (**Movie 2, Fig. 3B**). At initial aggregation stages, we observed that neighboring adherent cells contact each other by their filopodia and quickly coalesce into tight aggregates. The timescale of this initial cell-cell aggregation is too sudden to be explained by changes in gene expression. We therefore hypothesized that aggregation initiation does not require the expression of new proteins. To further test this hypothesis, we inhibited protein translation (using cycloheximide) and tested if these non-translating cells could still aggregate in response to induction with FBS (containing LDL and HDL). After confirming that cycloheximide inhibited translation in *Capsaspora* (**Fig. S5**), we found that cycloheximide *did not* inhibit aggregation (**Fig. 3C**). This finding demonstrates that changes in gene expression are not required for the initial lipid-induced aggregation of *Capsaspora*. Therefore, further investigation into the mechanism of *Capsaspora* aggregation will require post-translational analysis methods.

**Figure 3:**
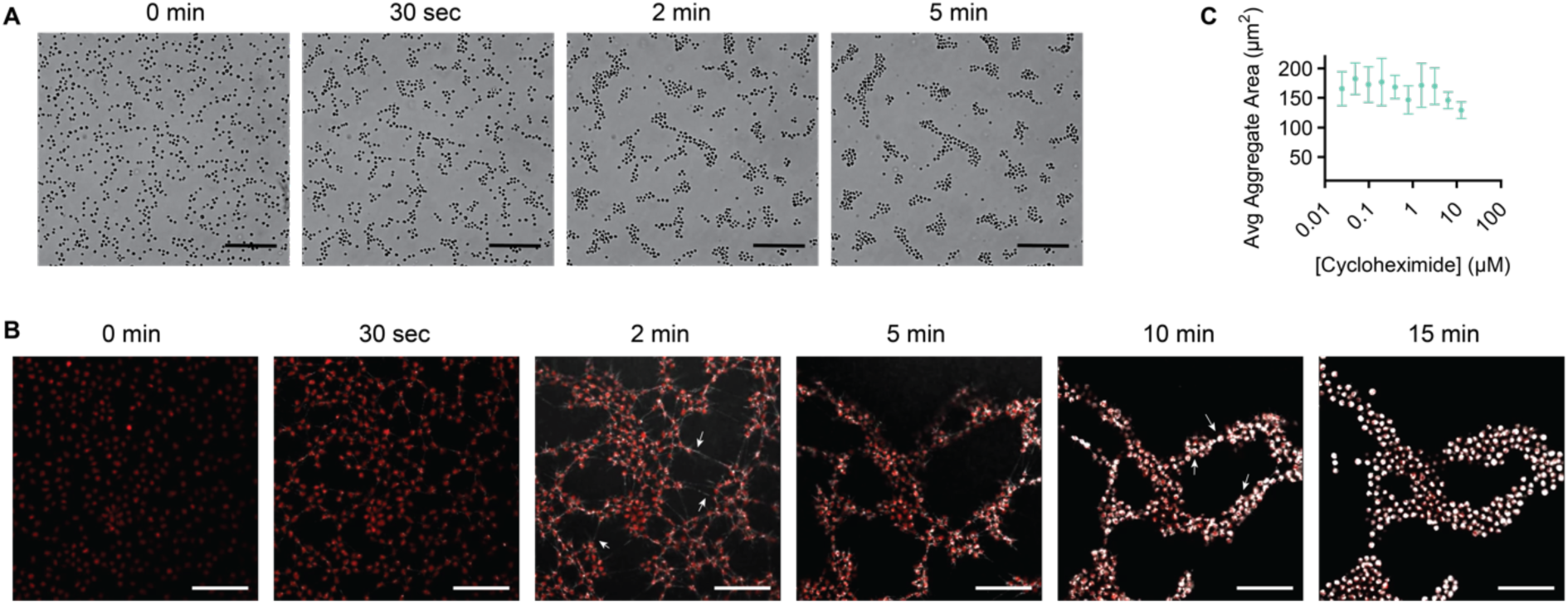
*Capsaspora* aggregation starts in seconds and does not require translation of new proteins. (A) Time series of brightfield images after induction by 5% (v/v) FBS. (B) Time series of fluorescent microscopy images after addition of fluorescent PC particles. Red coloring is TdTomato-labeled *Capsaspora* cells. White coloring is fluorescent PC particles (DOPC/TopFluorPC [20:1 (w/w)]). Fluorescent PCs are rapidly endocytosed, correlating with cells coalescing into aggregates. Arrows in the 2-minute panel show fluorescent PC particles along long filopodial connections between cells. Arrows in the 10-minute panel show fluorescent PC particles accumulate in puncta. (C) Average aggregate area after 30 minute pre-treatment with cycloheximide and induction with 5% (v/v) FBS. Error bars represent standard error of the mean (n=3).

## DISCUSSION

We discovered that *Capsaspora* aggregation is post-translationally induced by an assortment of zwitterionic diacyl phospholipids. We further found that the lipids are endocytosed, and clathrin-mediated endocytosis is necessary for the lipids to induce *Capsaspora* aggregation.

Although we previously observed that *Capsaspora* aggregation is induced by DOPC,^8^ here we discovered that *Capsaspora* aggregates in response to any tested phosphatidylcholine (PC) lipid with at least one unsaturation in at least one lipid tail. Sphingomyelins were also very active. This promiscuous activation of aggregation suggests that *Capsaspora* (or its ancestors) may have evolved to aggregate in response to nutrients that indicate the presence of a variety of prey (not only snail host environments). As an osmotrophic feeder, *Capsaspora* is believed to secrete digestive enzymes to release nutrients from prey. The aggregation of many cells around a common prey cell/colony/tissue maximizes the efficiency of nutrient acquisition by allowing the cells to share their enzymes and released nutrients as common goods at a high local density (**Fig. 4A**).^33^ Zwitterionic lipids are universally abundant in the membranes of animals (mostly PCs), bacteria (mostly PEs), fungi (PCs and PEs), and algae (PCs). Since these lipids are frequently released in extracellular vesicles by living cells^34^ and by lysing cells,^35^ they would be excellent cues to signal the presence of prey. Therefore, although DOPC and other PCs in snail serum may be an aggregation cue for *Capsaspora* in response to its snail host environment,^8^ this aggregation behavior may also be relevant outside of the *Capsaspora*-snail symbiosis. It could help *Capsaspora* to colonize yet-unknown animal hosts or form feeding aggregates around colonies of other prey. This lipid-activated aggregation behavior may have even evolved in the ancestors of *Capsaspora* to feed on unicellular prey in a pre-metazoan world. To test this hypothesis, work is currently underway in our laboratory to discern if other filastereans also aggregate in response to zwitterionic lipids.

**Figure 4:**
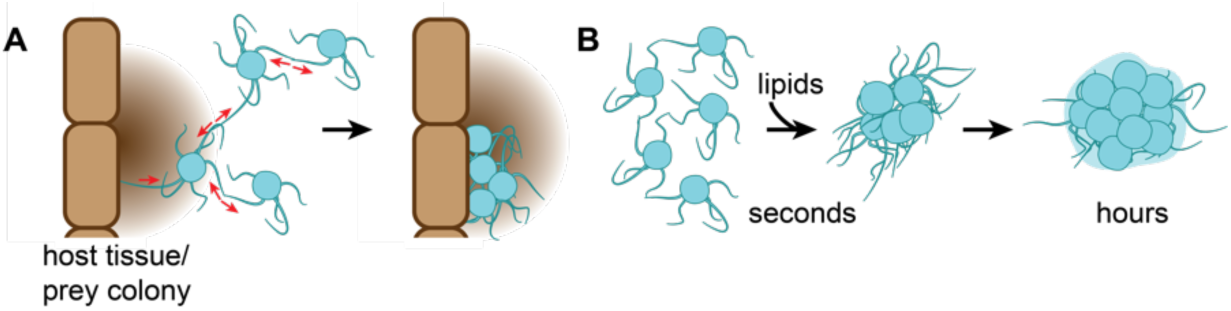
Generality and kinetics of *Capsaspora* aggregation. (A) Immediate cell-cell contraction leads to aggregation in response to zwitterionic lipids; this response may generally improve *Capsaspora* feeding efficiency by increasing cell density around any nutrient-rich host tissue or colony of prey cells. (B) *Capsaspora* aggregation likely involves stages of development; initial aggregation of cells is followed by changes in gene expression that may retract filopodia and produces extracellular matrix.^2^

In line with the nutritional potential of lipids, we have found that LDLs and PC vesicles are endocytosed by *Capsaspora*. The aggregation response appears dependent on the clathrin-mediated endocytosis of these lipids, perhaps indicating that a lipid-sensing receptor signals from within endocytic vesicles.^36^ The convergence of lipid particle endocytosis and cellular signaling is reminiscent of lipoproteins and their receptors in animals.^37–39^ Several proteins related to the low density lipoprotein receptor (LDLR) are responsible for lipoprotein uptake and/or cellular signaling processes. *Capsaspora* possesses transmembrane proteins that harbor the cytosolic NPXY motif, which classically drives endocytosis of LDLR homologs and some signaling processes.^38^ The large repetitive structure of lipid vesicles may also prove ideal for the multivalent interactions that assemble receptor multimers at the cell surface that typically activate endocytosis and/or signaling from LDLR homologs.^38^ Work is underway to identify the receptor(s) in *Capsaspora*.

The rapid generation of aggregates via post-translationally induced cell-cell compaction provides new insight into the early stages of *Capsaspora*’s aggregation process. The initial step of aggregation is clearly “activation” of proteins already present in the cells—not increased production of adhesion proteins and extracellular matrix material, as was previously suspected following gene expression analysis of mature aggregates.^1, 2^ However, the previously reported changes in gene expression likely do arise as aggregates mature, especially in the presence of agitation.^2^ Therefore, these data suggest multiple stages of aggregate maturation (**Fig. 4B**). An initial cell-cell contraction event rapidly assembles cells in a tight colony within a few minutes. Subsequent gene expression changes further mature the colony, which may include filopodial retraction and production of extracellular matrix among other physiological changes.^2^ A comprehensive time-series analysis of gene expression changes will further inform the full progress of aggregate maturation.

## CONCLUSION

In sum, we discovered that *Capsaspora* aggregation is induced by a wide panel of abundant zwitterionic diacyl phospholipids. The diverse nature of the inducing lipids suggests that the aggregation response might be general to many nutrient-rich environments (not just host snails) and may enable *Capsaspora* (or its ancestors) to efficiently prey on other organisms. Moreover, *Capsaspora* lipid-induced aggregation utilizes clathrin-mediated endocytosis of lipids. This aggregative process occurs remarkably fast (within seconds) via post-translational processes. The sudden cell-cell compaction observed in early aggregation reveals that the process of aggregate development is more complex than previously appreciated, requiring further analysis for comparison with similar cell-cell adhesion and contraction behaviors in multicellular animals. Overall, these insights advance our understanding of *Capsaspora* cellular aggregation, which is a potential key to the unicellular-to-multicellular transition in the animal lineage and may also be an essential behavior to support an anti-schistosome symbiosis.

## MATERIALS AND METHODS

### Cell strain and growth conditions

*Capsaspora owczarzaki* cell cultures (strain ATCC®30864) were grown axenically in 25 cm^2^ culture flasks with 6 mL ATCC media 1034 (modified PYNFH medium: 10 g/L peptone, 10 g/L yeast extract, 1 g/L yeast ribonucleic acid, 15 mg/L folic acid, 1 µg/mL hemin, 2.66 mM KH_2_PO_4_, and 3.52 mM Na_2_HPO_4_ at pH 6.5 in water) containing 10% (v/v) heat-inactivated Fetal Bovine Serum (FBS), hereafter *growth media*, in a 23°C incubator. Adherent stage cells (filopodiated amoebae) at the exponential growth phase were obtained by passaging ~100–150 µL of adherent cells at ~90% confluence in 6 mL of growth media and grown for 24–48 hours at 23°C until ~100% confluent.

### General aggregation assay methods

All aggregation assays were performed at room temperature. Brightfield imaging was performed using the following instruments: Leica DMi1 inverted microscope with an MC120 HD camera, Leica DMIL inverted microscope with Flexacam C3 camera, an Olympus OSR spinning disk confocal microscope with a Hammamatsu Flash 4 V2 camera, and an Incucyte S3 Live-Cell Analysis System, and an A1 Nikon Scanning Confocal with a Hammamatsu Orca-Flash 4.0 sCMOS camera. Depending on well size and microscope used, each well was imaged at up to 3 distinct locations using 5X or 10X magnification. Average aggregate areas were typically measured by batch processing with a standard macro script in Fiji Imaging Software^7, 40^ (see *Image analysis* below).

#### Aggregation assay on ultra-low attachment plates

Two days before the assay, 100% confluent adherent cells growing in 25 cm^2^ culture flasks were given fresh growth media (termed the “feed step”). One day before the assay, cells were washed and resuspended in FBS-free assay media and allowed to sit overnight (termed the “starve step”). After starvation, the day of the assay, 8*10^5^ cells were seeded in 180 µL of FBS-free media per well in a 96-well ultra-low attachment microplate (#CLS3474, Corning) and allowed to settle for 2 hours. Putative aggregation inducers were added such that the total volume in a well was 200 µL. Typically, aggregates were assessed by microscopy after 90 min.

#### Image analysis for aggregation assays

Average aggregate areas were typically measured by batch processing with a standard macro script in Fiji Imaging Software version 2.1.0/1.53c.^7, 40^ Briefly, the macro steps included: set the scale of the image appropriate for the microscope conditions, convert the image to binary, analyze particles (size 0-infinity), export results to clipboard. A copy of the FIJI macro is available upon request.

### Structure activity relationship studies (related to Figure 1 and Figure S1 and S2)

The following synthetic lipids were acquired from Avanti Polar Lipids unless otherwise specified:

| **Lipid name** | **abbreviation** | **catalog number** |
| --- | --- | --- |
| 1-palmitoyl-2-oleoyl-glycero-3-phosphocholine | POPC | 850457C |
| 1,2-dioleoyl-sn-glycero-3-phosphocholine | DOPC | 850375P |
| 3-palmitoyl-2-oleoyl-sn-glycero-1-phosphocholine | Ent-POPC | 850855C |
| 1,2-di-O-(9Z-octadecenyl)-sn-glycero-3-phosphocholine | diether-PC | 999989P |
| 2-((2,3-bis(oleoyloxy)propyl)dimethylammonio)ethyl ethyl phosphate | DOCPe | 850312P |
| 1-palmitoyl-2-oleoyl-sn-glycero-3-ethylphosphocholine (chloride salt) | EPC | 890705P |
| N-oleoyl-D-erythro-sphingosylphosphorylcholine | 18:1 SM | 860587C |
| N-palmitoyl-D-erythro-sphingosylphosphorylcholine | 16:0 SM | 860584P |
| 1-palmitoyl-2-oleoyl-sn-glycero-3-phosphate (sodium salt) | POPA | 840857C |
| 1-palmitoyl-2-oleoyl-sn-glycero-3-phosphoethanolamine | POPE | 850757P |
| Phosphocholine chloride calcium salt tetrahydrate | phosphocholine | sigma P0378 |
| Choline chloride | choline | Fisher A1582822 |
| 1-palmitoyl-2-hydroxy-sn-glycero-3-phosphocholine | 16:0 LPC | 855675P |
| 1-oleoyl-2-hydroxy-sn-glycero-3-phosphocholine | 18:1 LPC | 845875P |
| 1,2-dimyristoleoyl-sn-glycero-3-phosphocholine | 14:1 PC | 850346C |
| 1,2-dipalmitoleoyl-sn-glycero-3-phosphocholine | 16:1 PC | 850358C |
| 1,2-dilinolenoyl-sn-glycero-3-phosphocholine | 18:3 PC | 850395C |
| 1,2-dieicosenoyl-sn-glycero-3-phosphocholine | 20:1 PC | 850396C |
| 1,2-diarachidonoyl-sn-glycero-3-phosphocholine | 20:4 PC | 850397C |
| 1,2-dimyristoyl-sn-glycero-3-phosphocholine | DMPC | 850345P |
| 1,2-dipalmitoyl-sn-glycero-3-phosphocholine | DPPC | 850355C |
| 1,2-distearoyl-sn-glycero-3-phosphocholine | 18:0 PC | 850365C |
| 1,2-di[(8Z)octadecenoyl]-sn-glycero-3-phosphocholine | 18:1 (8cis) PC | 792486C |
| 1,2-dielaidoyl-sn-glycero-3-phosphocholine | 18:1 (9trans) PC | 850376C |
| 1,2-divaccenoyl -sn-glycero-3-phosphocholine | 18:1 (11cis) PC | 790626C |
| 1,2-dinervonoyl-sn-glycero-3-phosphocholine | 24:1 PC | 850399C |

#### Preparing lipid vesicles from pure lipid samples (related to Fig. 1 and Fig. S1 and S2)

Lipids commercially obtained (pure synthetic lipids) were dissolved in chloroform in glass vials. 0.2mg was transferred to a 1.7 mL microcentrifuge tube or a 1 dram glass vial. The chloroform was evaporated using a gentle stream of nitrogen and the resulting lipid film was then resuspended in 1 mL each of 1X PBS or FBS-free assay media. The tubes were vortexed for 30 seconds before sonication. Lipid solutions were sonicated on ice with a 50% duty cycle at medium setting for 10 minutes using a single probe attached to a Branson Sonifier Cell-Disruptor 185 or in sets of 4 using a 4 probe sonicator horn attached to a Fisherbrand Model 505 Sonic Dismembrator (FB505A110). During sonication, tubes were kept on ice and tube bottoms were set about 3 mm from the sonicator tip. Lipid solutions went from slightly cloudy to clear after sonication. Sonicated lipids were stored at 4°C for no more than 2 days before use.

Crude lipids were concentrated 10X using a >30 kDa cutoff filter immediately before addition into aggregation assay wells. The concentration of lipids were reported in µg/mL based on mass of lipid/volume of the assay well.

#### Testing lipid vesicles for aggregation (related to Fig. 1 and Fig. S1 and S2)

Sonicated lipids were concentrated 10X using an Amicon Ultra 30 kDa cutoff filter and diluted with 1X PBS to the desired concentration before testing. The standard aggregation assay on ultra-low attachment plates was used to test for aggregation induction. Aggregates were induced using the desired lipid concentration (calculated in µg/mL of lipid) and 5% (v/v) of 1X PBS or sonicated FBS-free assay media was used as a negative control depending on what the lipids were prepared in. Aggregates in triplicate wells were imaged every 30 minutes and analysis was performed on images from T-90 minutes. Average aggregate areas were measured using a macro in FIJI as described above.

#### Transmission Electron Microscope (TEM) imaging of prepared lipid vesicles (related to Fig. S1)

Before preparation of samples, Formvar/Carbon film 300 mesh Nickel Grids (Electron Microscopy Sciences FCF300-Ni-25) were ionized using a Pelco easiGlow Discharge System at 0.3 mbar and 15 mAmps for 2 minutes. 10 µL of prepared lipid samples were added to the ionized grids and allowed to sit for 5 minutes. Excess sample was removed from the grids by gently touching Whatman filter paper (Sigma, WHA1001329) to the edge. Immediately after removing samples (before grid dries) 10 µL of 2% (v/v) Uranyl Acetate pre-diluted in water was added to grids and allowed to sit for 10 seconds. Then the stain was removed with the edge of Whatman filter paper. Grids were allowed to dry completely before imaging (about 10 minutes). Prepared grids were imaged immediately after preparation using a JEOL-JEM 1010 transmission electron microscope (TEM) with an 80 kV operating voltage, equipped with a 1k x 1k Gatan CCD camera (MegaScan model 794) using a tungsten filament as its electron source.

#### Mass spectrometric analysis of oxidized lipids in prepared vesicle samples (Fig. S3)

For analysis of the amount of oxPC in prepared lipid vesicle samples, 1 mL of vesicles prepared and concentrated to 10x as described above (or a control with 0.2 mg of lipid from a stock vial never exposed to sonication) was resuspended in 0.5 mL of 2M NaCl in water and extracted using 1.9 mL of 1:2 chloroform/methanol for 15 minutes at room temperature. Then an additional 0.6mL of water and 0.6mL of chloroform was added and vortexed and the sample was allowed to separate. The lower layer (organic) was collected, and the upper layer was extracted with and additional 0.8mL of chloroform. The organic layers were combined and the solvent was evaporated using a Savant SpeedVac SPD2030 at 45°C under 5.1 millitorr of vacuum pressure for 4 hours. The dried lipids were resuspended in 75 µL of 2:1 isopropanol/methanol. Samples were centrifuged briefly to remove any suspended particles. High-resolution electrospray ionization (HR-ESI) mass spectra with collision-induced dissociation (CID) MS/MS were obtained using an Agilent LC-q-TOF mass spectrometer 6530 equipped with an Agilent 1290 uHPLC system. Metabolites were separated using a Luna 5 μm C5 100 Å LC column (Phenomenex 00D-4043-E0). Mobile phase A was 95% (v/v) Water, 5% (v/v) Methanol, 0.1% (v/v) Formic Acid, and 5 mM Ammonium Formate. Mobile phase B was 60% (v/v) Isopropanol, 35% (v/v) Methanol, 5% (v/v) Water, 0.1% (v/v) Formic Acid, and 5 mM Ammonium Formate. After initially holding 0% phase B for 5 min at 0.1 mL/min, a linear gradient from 20% phase B to 100% phase B was applied over 40 min with a flow of 0.4 mL/min before holding at 100% phase B for another 5 min with a flow of 0.5 mL/min. Data-dependent acquisition was employed to fragment the top masses in each scan. Collision-induced dissociation was applied using a linear formula that applied a higher voltage for larger molecules (CID voltage = 10 + 0.02 m/z) for metabolite profiling and identification. Mass traces collected in positive mode were analyzed in MassLynx software v4.1 and the Extracted ion chromatogram is of m/z 650.439 ± 15 ppm.

### Endocytosis Studies (Related to Figure 2 and S4)

#### Uptake of BODIPY labeled fluorescent LDL (related to Fig. 2B)

Cells were prepared as if for a normal aggregation assay as described above. Cells were seeded in assay media in ibidi chambered glass microscope slides (Ibidi 80827) and allowed to settle for 2 hours. Assay media was replaced with Chernin’s balanced salt solution (CBSS+, 2.8 g/L sodium chloride, 0.15 g/L potassium chloride, 0.07 g/L sodium phosphate dibasic, 0.45 g/L magnesium sulfate heptahydrate, 0.53 g/L calcium chloride dihydrate, 0.05 g/L sodium bicarbonate, 1 g/L glucose, and 1 g/L trehalose at pH 7.2 in water)^41^ buffer for imaging by sequential aspiration and replacement until liquid in wells was clear (about 3 times) and the total volume in the well was 400 µL. Storage buffer for fluorescent low-density lipoprotein (BODIPY LDL, Fisher L3483) was replaced with CBSS+ buffer before adding. 80 µg/mL of BODIPY LDL was added to the microscope slide and allowed to incubate for 30 minutes, then 5 µg/mL of Hoechst DNA stain (Fisher BDB561908) was added and allowed to sit for 5 minutes before removing and replacing with fresh buffer immediately before imaging on an OMX-SR 3D-SIM system with an excitation laser of 488nm for BODIPY LDL and 405nm for Hoechst at a magnification of 60x. Images are example slides with a scale bar of 5 µm.

#### Uptake of fluorescent PC particles (related to Fig. 2C)

Cells were prepared as if for a normal aggregation assay as described above. Fluorescent NBD-PC (Avanti 810130C) was doped in at a ratio of 20 POPC to 1 NBD-PC before making vesicles via sonication as described above. Cells were seeded in FBS-free assay media into wells in a 96-well black walled flat bottom plate (Corning 353219) and allowed to settle for 2 hours. Assay media was replaced with Chernin’s balanced salt solution (CBSS+) buffer for imaging by sequential aspiration and replacement until liquid in wells was clear (about 3 times) and the total volume in the well was 200 µL. 100 µg/mL of NBD-PC:POPC particles were added to cells and allowed to incubate for 30 minutes before imaging on an Olympus spinning disk confocal with 488nm excitation laser with a magnification of 20x. Images shown are example wells with a scale bar of 50 µm.

#### Inhibition of uptake of pHrodo LDL particles using Dynole 34-2 (related to figure 2D–E)

Cells were prepared for a normal aggregation assay as described above. Cells were seeded in FBS-free assay media into ultra-low attachment assay plates and allowed to settle for 2 hours. A dilution series of Dynole 34-2 (Neta scientific, CAYM-34073-1) in DMSO was prepared and added to assay wells in triplicate. In all cases, DMSO concentrations were kept constant at 0.5% in assay wells except for one control with no DMSO and one control with no pHrodo LDL. Cells were incubated with Dynole 34-2 for 30 minutes. The storage buffer for pHrodo LDL (Fisher, I34360) was replaced with PBS buffer immediately before using in the assay. 80 µg/mL of pHrodo LDL was added to each well and allowed to sit for 30 minutes. Cells were then fixed using 4% formaldehyde and assay media was replaced with CBSS+ for imaging. 5 µg/mL of Hoechst DNA stain (Fisher, BDB561908) was added and allowed to sit for 5 minutes before removing and replacing with fresh CBSS+ buffer. Cells were imaged on an Olympus microscope with widefield CFP and RFP lamps at 20x magnification. Images shown are example images of cells in wells with DMSO but no Dynole 34-2. Images were quantified in batch using Fiji Imaging Software version 2.1.0/1.53c. Briefly the macro was set to find edges, convert to binary, despeckle, and count particles in the Hoechst channel and in the pHrodo channel. Plot is the percent of Hoescht stained cells that were also stained with pHrodo, final result was analysis of duplicate images of triplicate wells (n=6).

#### Inhibition of *Capsaspora* aggregation using chemical inhibitors of endocytosis (related to Fig. 2F–G and Fig. S4)

Normal aggregation assays were set up according to the protocol above. Inhibitors were purchased and prepared according to solubility. In cases where inhibitors were soluble in DMSO, the final DMSO concentration in assay wells was kept at a constant 0.5% and a DMSO control was included. In cases where inhibitors were soluble in ethanol, the final concentration of ethanol in assay wells was kept at a constant 1% and an ethanol control was included. After allowing cells to settle in assay plates, cells were treated with chosen inhibitors for 30 minutes before inducing aggregation with 5% (vol/vol) FBS. Assays were performed in triplicate and aggregates were analyzed 90 minutes after induction. A list of inhibitors is available as supplementary Table below.

| **Inhibitor** | **Source** | **Catalog number** | **Solute** |
| --- | --- | --- | --- |
| Dynole 34-2 | Neta Scientific | CAYM-34073-1 | DMSO |
| Pitstop II | Abcam | ab120687 | DMSO |
| Dynasore | Neta Scientific | CAYM-14062-5 | DMSO |
| MiTMAB | Abcam | ab120468 | DMSO |
| OctMAB | Abcam | ab120468 | DMSO |
| CK-666 | Sigma | 182515-25MG | DMSO |
| Chlorpromazine | Sigma | C8138-5G | DMSO |
| Ponesimod | Sigma | ADV638391874-25MG | DMSO |
| ABMA | Med Chem Express | HY-124801 | DMSO |
| EGA | Sigma | 5093060001 | DMSO |
| Chloroquine | Sigma | C6628-25G | DMSO |
| Monensin | Sigma | M5273-500MG | DMSO |
| Cycloheximide | Sigma | 01810-1G | Water |

#### Videos of *Capsaspora* aggregation (related to Fig. 3A and Movie 1)

*Capsaspora* cells were prepared for a standard aggregation assay as described above. Cells were allowed to settle in ultra-low attachment plates for 2 hours. Plates were placed on Leica DMi1 inverted microscope and focused on 10x magnification with brightfield illumination. Video recording was started and 5% (vol/vol) FBS was added gently to an edge of the assay well about 10 seconds after recording had started and cells were recorded for 5 minutes continuously. Video was converted to black and white to show cells more clearly and compressed from its original size before upload using ffmpeg tool build 20200213-6d37ca8.^42^

#### Timelapse imaging of *Capsaspora* uptake of fluorescent PC particles (related to Fig. 3B and Movie 2)

Capsaspora cells constitutively expressing TdTomato fluorescent protein^8^ were prepared as if for a standard aggregation assay as described above. About 1.6*10^6^ cells were added in FBS-free assay media to an ibidi chambered glass microscope slide and allowed to settle for 2 hours. Directly before imaging, assay media was removed and replaced with CBSS+ buffer to reduce background fluorescence. Fluorescent PC particles were made by combining DOPC (Avanti Polar Lipids 850375P) and TopFluorPC (Avanti Polar Lipids 810281C-1mg) in a ratio of 20 DOPC to 1 TopFluorPC (w/w). Lipids vesicles were prepared by mixing 2mg of total lipids dissolved in chloroform in a glass vial, evaporating the chloroform with a gentle stream of nitrogen to form a thin film, and them resuspending lipids in 1mL of PBS buffer. Lipids were then extruded through 100 nm pores in an extruder (Avanti Polar Lipids 610000). Lipids were used in the assay the same day. Cells were placed on an Olympus spinning disk microscope and images were set up to be taken of each channel (561 nm excitation for TdTomato and 488 nm excitation for TopFluorPC) and imaging was started. After 2 imaging cycles (~12 seconds) fluorescent PCs were added gently to one corner of the well while the microscope recording was still running. Images were taken every 5.6 seconds for ~15 minutes. Stacks of images were compressed and converted into videos using Fiji software.

#### Inhibition of new protein synthesis using Cycloheximide (related to Fig. 3C and Fig. S5)

Normal aggregation assays were set up according to the protocol above. A dilution series of the protein synthesis inhibitor cycloheximide (Sigma 01810) was created in water. After allowing cells to settle in assay plates for 2 hours, cells were treated with cycloheximide inhibitors for 30 minutes before inducing aggregation with 5% (v/v) FBS. Images for aggregation assay were assessed 90 minutes after addition of FBS.

#### Measuring new protein synthesis using incorporation of radioactive amino acid [^35^S] methionine (related to Fig. S5)

Cells were prepared as if for a standard aggregation assay as described above. About 480 million cells were pelleted with centrifugation at 1000 x g and resuspended into 12 mL of FBS-free assay media. These cells were divided into 24 portions, each with 0.5 mL of cells (about 20 million cells) in 1.7 mL centrifuge tubes. Cycloheximide dilutions were prepared in water and added to cells to reach the desired final concentration. A negative control with water and positive control with 4% formaldehyde were also prepared. Cells were treated with cycloheximide (or formaldehyde) for 30 minutes. Then, 20.4 µCi of [^35^S] L-methionine (Revvity/Perkin Elmer NEG009A500UC) was added to each tube and allowed to incubate for 2 hours. Cells were then pelleted by centrifugation and resuspended in 0.5 mL of cold cell lysis buffer (NP-40 buffer + 1 mg/mL SDS + 5 mg/mL sodium deoxycholic acid + 0.38 mg/mL EGTA). Cells were incubated on ice for 30 minutes and then pipetted vigorously before centrifugation at 15,000 x g for 10 minutes to remove debris. The supernatant was then mixed 1:1 with TCA (acetone + 20% trichloroacetic acid + 0.07% 2-mercaptoethanol) and allowed to sit for 45 minutes on ice to precipitate proteins. After precipitation, samples were centrifuged at 15,000 x g for 10 minutes and the pellet was washed with 1 mL of acetone before air drying. The dry pellets were then resuspended in 40 µL of PBS buffer and the whole sample was spotted onto whatman filter paper. The paper was allowed to dry completely before covering with plastic wrap and placing in a phosphor plate chamber overnight. The next day, the phosphor plate was imaged using a Typhoon 9210 Variable Mode Imaging System. Quantification of the intensity of the dot blot was performed using Fiji image software by measuring the average intensity of each spot.

## AUTHOR CONTRIBUTIONS

Conceptualization, R.Q.K., J.P.G.; Methodology, R.Q.K., J.P.G.; Investigation, R.Q.K., E.B.G., H.J.R., L.P.B., A.M.G. N.R.R.; Writing – Original Draft, R.Q.K., J.P.G.; Writing – Review & Editing, R.Q.K., N.R.R., J.P.G.; Visualization – R.Q.K., J.P.G.; Supervision, J.P.G.; Funding Acquisition, J.P.G.

## MATERIALS & CORRESPONDENCE

Correspondence and material requests should be addressed to J. P. Gerdt.

## ACKNOWLEDGMENTS

We thank the Light Microscopy Center at Indiana University for support in image acquisition and analysis (funding provided by the NIH grant NIH1S10OD024988-01). We also thank the Indiana University Nanoscale Characterization Facility, Electron Microscopy Center, Laboratory for Biological Mass Spectrometry, and Physical Biochemistry Instrumentation Facility for use of their instruments. We also thank the entire Gerdt lab for insights and support that helped advance this project. This work was supported by the National Institutes of Health (R35GM138376) to J.P.G. R.Q.K. was supported by an NIH training grant (T32GM131994). The content of this paper is solely the responsibility of the authors and does not necessarily represent the official views of the National Institutes of Health.

## Supplementary Information

### SUPPLEMENTARY FIGURES

**Figure S1:**
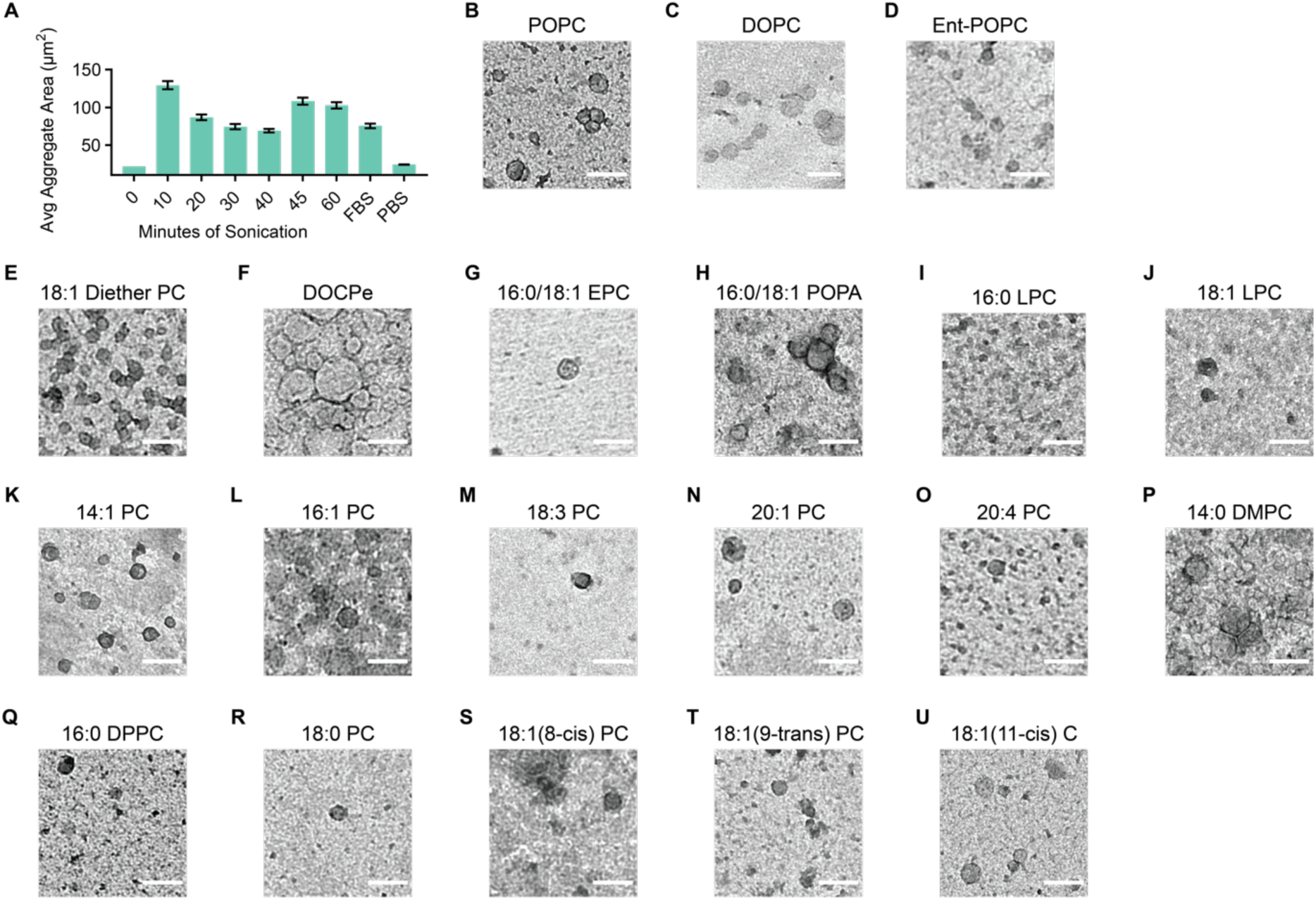
Lipoproteins, Sonication, and TEM images for Structure-Activity Relationship Study. (A) Average aggregate area resulting from addition of POPC prepared with varying sonication times. Preparation of PC lipids into vesicles by sonication is necessary for aggregation induction; adding lipids without sonication failed to induce aggregation. (B-U) Transmission electron microscope (TEM) images of prepared lipid vesicles stained with 1% uranyl acetate. Scale bars are 100 nm.

**Figure S2:**
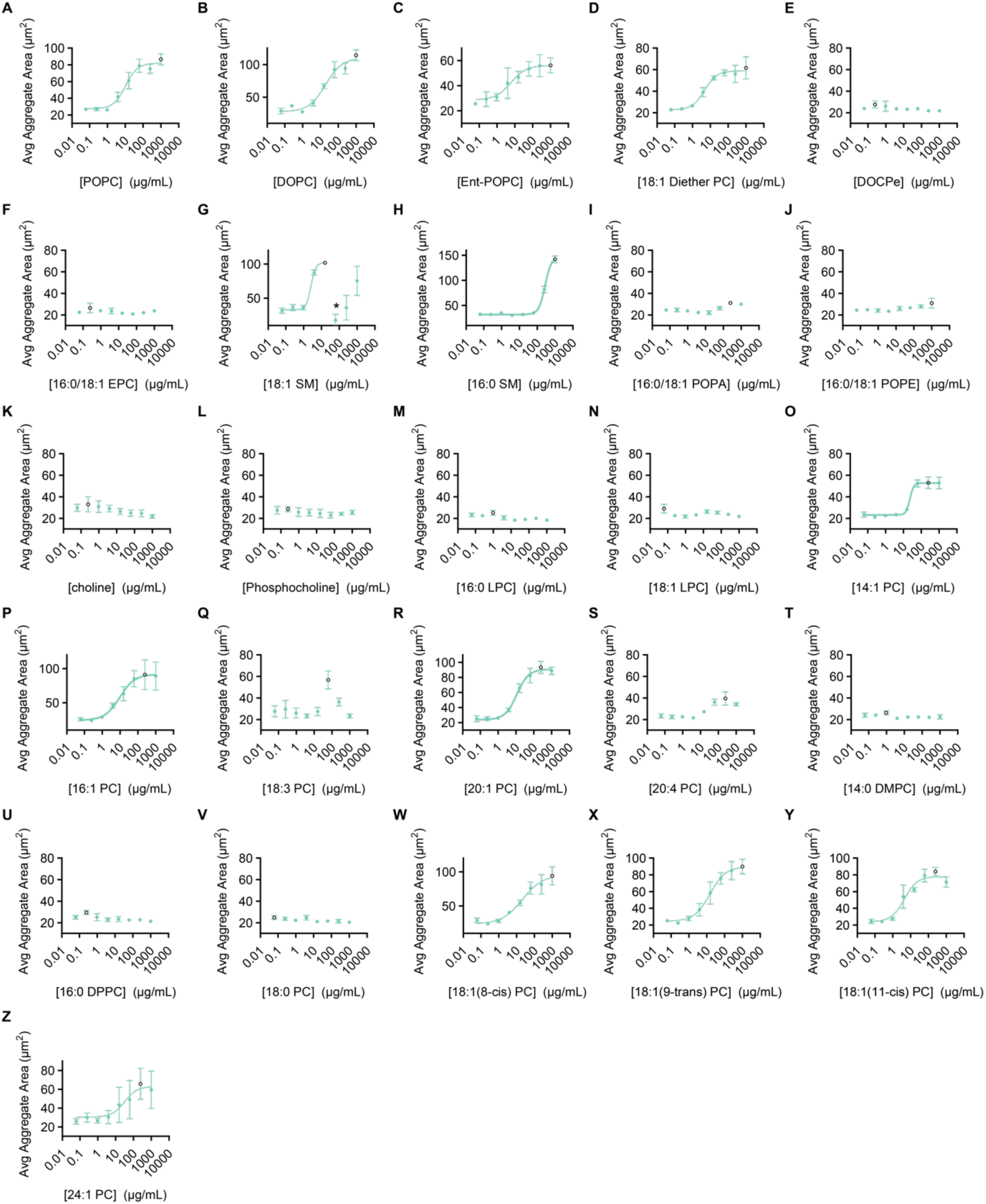
Individual Aggregation Curves for Structure Activity Relationship Study. (A-Z) Individual dose response curves for all tested lipids (the black circle represents the largest aggregate value, which was the selected concentration reported in the main text). All error bars represent SEM of biological triplicates (n=3).

**Figure S3:**
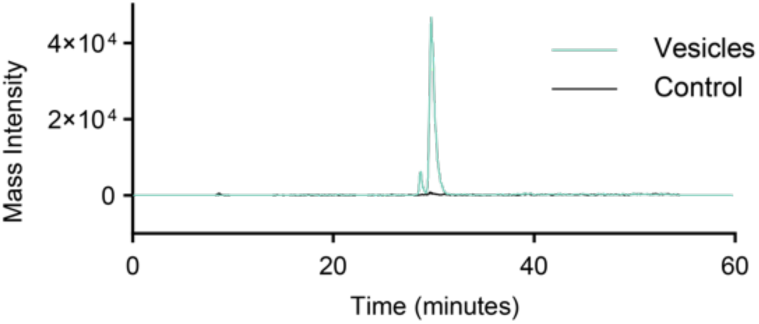
Liquid chromatography mass spectrometry (LCMS) analysis of prepared POPC lipid vesicles indicates a low abundance of oxidized PC (oxPC) present. Extracted ion chromatogram is of m/z 650.439 ± 15 ppm. The oxPC peak is ~1000x more intense after sonication into vesicles.

**Figure S4:**
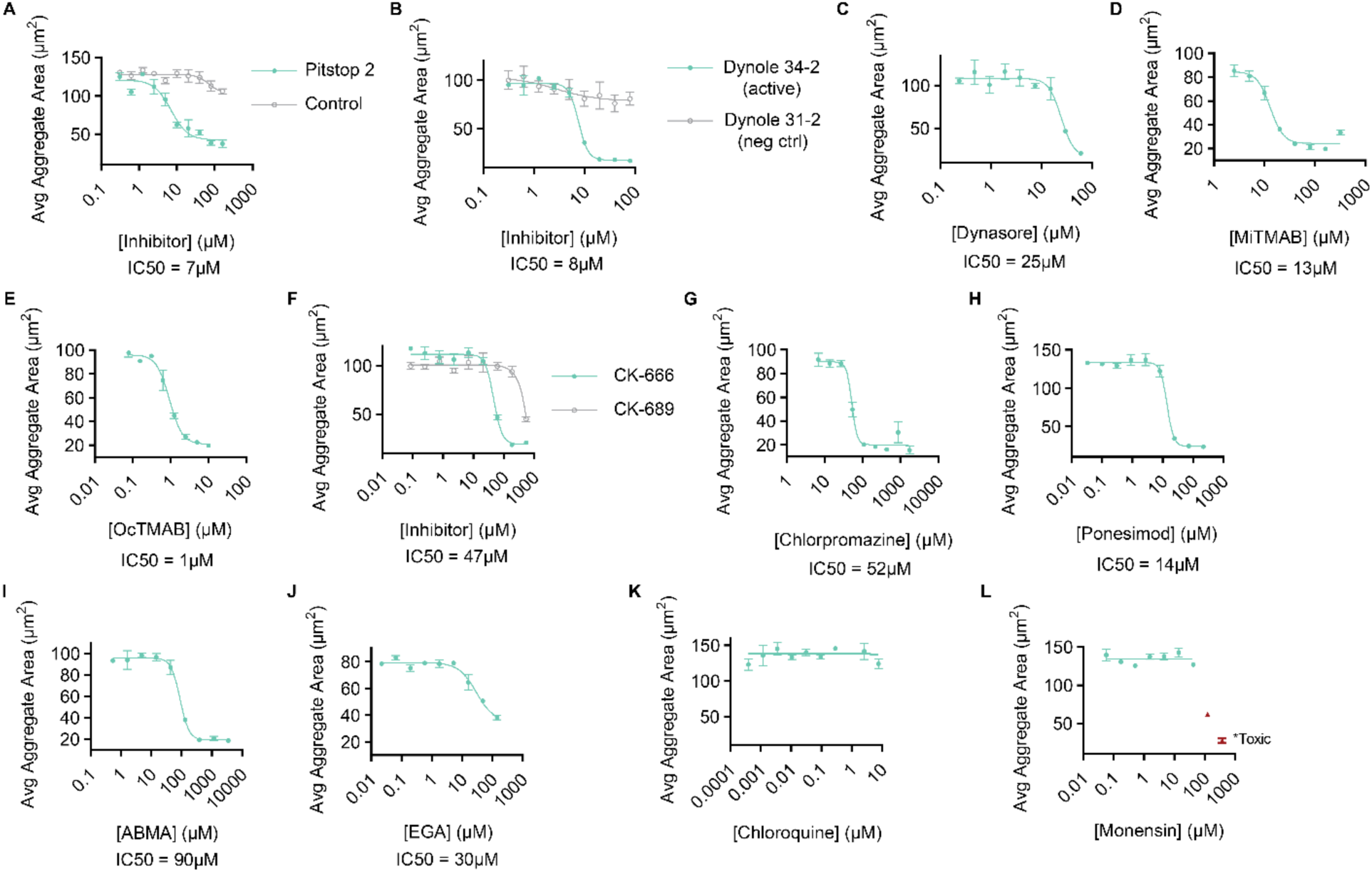
Individual dose response curves for all endocytosis inhibitors. (A)-(L) Average aggregate areas resulting from a dilution series of various endocytosis inhibitors. Aggregation was induced with 5% (v/v) FBS after 30 minutes of treatment with each inhibitor. (A) The clathrin recruitment inhibitor, Pitstop 2 inhibited aggregation while it’s negative control molecule did not. The IC50 of Pitstop in Capsaspora was found to be 7 µM. The reported IC50 for Pitstop is reported to be 12 µM in HeLa cells.^1^ (B) The dynamin inhibitor Dynole 34-2^2^ inhibited aggregation while it’s negative control molecule Dynole 31-2 did not. (C) The dynamin inhibitor Dynasore^3^ inhibited aggregation although not as strongly as Dynole 34-2. (D/E) The dynamin recruitment inhibitors MiTMAB and OcTMAB^4^ strongly inhibited aggregation. (F) The actin polymerization inhibitor CK-666^5^ prevented aggregation. (G) The clathrin decoating inhibitor chlorpromazine^6^ inhibited aggregation. (H) The endosome maturation inhibitor Ponesimod^7^ inhibited aggregation. (I) The endosome maturation inhibitor ABMA^8^ inhibited aggregation. (J) EGA even at the highest concentration of solubility still had many visible aggregates (although looser) and so was considered not active. The IC50 of EGA is reported to be 1µM in A549 cells.^9^ (K) The lysosome acidification inhibitor chloroquine^10^ did not inhibit aggregation. (L) The lysosome pH acidification inhibitor Monensin^10^ did not inhibit aggregation.

**Figure S5:**
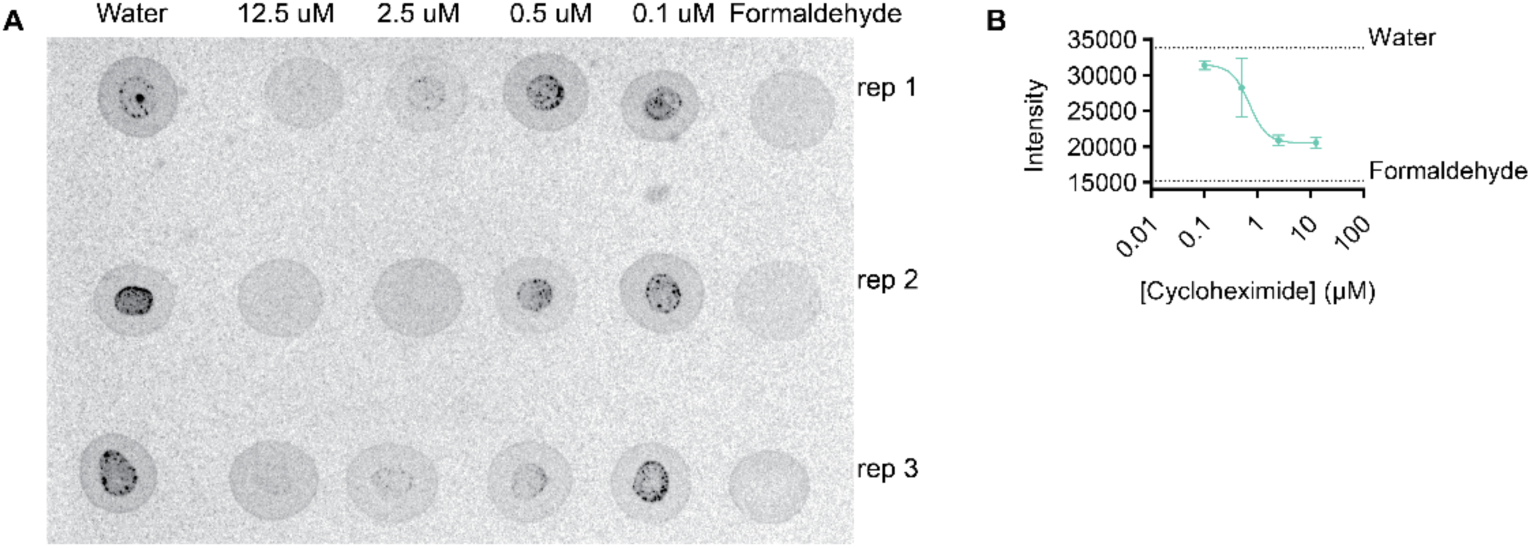
(A) phosphor plate image of incorporation of radioactive [^35^S] methionine in cycloheximide treated cells. Treated cells show no incorporation of amino acid at 12 µM cycloheximide, indicating no new proteins are being translated. (C) Quantification plot of phosphor plate results. Error bars are SEM (n=3).

### LEGENDS FOR MOVIES

**Movie 1: Aggregation of *Capsaspora* is fast.** Brightfield microscopy video of *Capsaspora* cells aggregating after induction with 5% (v/v) FBS. Continuous video taken with a MC120 HD camera. Original video was converted to a stack of frames for compression using FFmpegTool and frames from Movie 1 were used to generate the images in **Fig. 3A**. Scale bar is 50 µm and time in minutes:seconds is displayed in the top left corner. Time 00:00 corresponds to the addition of FBS.

**Movie 2: Aggregation of *Capsaspora* coincides with uptake of fluorescent PC particles.** Confocal microscopy video of *Capsaspora* cells expressing TdTomato (red) aggregating upon addition of 100 µg/mL of fluorescent PC particles (white). Particles localize first to filopodia and are ultimately internalized in puncta inside cells by 15 minutes. Video generated by taking images every 4.6 seconds for 15 minutes. Frames from Movie 2 were used to generate the images in **Fig. 3B**. Scale bar is 50 µm and time in minutes:seconds is displayed on the top left corner. Time 00:00 corresponds to the addition of fluorescent PCs.

## REFERENCES

1. Sebé-Pedrós A, Ballaré C, Parra-Acero H, Chiva C, Tena JJ, Sabidó E, Gómez-Skarmeta JL, Di Croce L, Ruiz-Trillo I. The Dynamic Regulatory Genome of Capsaspora and the Origin of Animal Multicellularity. Cell. 2016;165(5):1224–37; doi: 10.1016/j.cell.2016.03.034.

2. Suga H, Chen Z, de Mendoza A, Sebé-Pedrós A, Brown MW, Kramer E, Carr M, Kerner P, Vervoort M, Sánchez-Pons N, Torruella G, Derelle R, Manning G, Lang BF, Russ C, Haas BJ, Roger AJ, Nusbaum C, Ruiz-Trillo I. The Capsaspora genome reveals a complex unicellular prehistory of animals. Nat Commun. 2013;4:2325; doi: 10.1038/ncomms3325.

3. Stibbs HH, Owczarzak A, Bayne CJ, DeWan P. Schistosome sporocyst-killing Amoebae isolated from Biomphalaria glabrata. J Invertebr Pathol. 1979;33(2):159–70; doi: 10.1016/0022-2011(79)90149-6.

4. Hertel LA, Bayne CJ, Loker ES. The symbiont Capsaspora owczarzaki, nov. gen. nov. sp., isolated from three strains of the pulmonate snail Biomphalaria glabrata is related to members of the Mesomycetozoea. International Journal for Parasitology. 2002;32(9):1183–91; doi: 10.1016/S0020-7519(02)00066-8.

5. Owczarzak A, Stibbs HH, Bayne CJ. The destruction of Schistosoma mansoni mother sporocysts in vitro by amoebae isolated from Biomphalaria glabrata: an ultrastructural study. J Invertebr Pathol. 1980;35(1):26–33; doi: 10.1016/0022-2011(80)90079-8.

6. Ferrer-Bonet M, Ruiz-Trillo I. Capsaspora owczarzaki. Current Biology. 2017;27(17):R829–R30; doi: 10.1016/j.cub.2017.05.074.

7. Ros-Rocher N, Kidner Ria Q, Gerdt C, Davidson WS, Ruiz-Trillo I, Gerdt Joseph P. Chemical factors induce aggregative multicellularity in a close unicellular relative of animals. Proceedings of the National Academy of Sciences. 2023;120(18):e2216668120; doi: doi:10.1073/pnas.2216668120.

8. Kidner RQ, Goldstone EB, Laidemitt MR, Sanchez MC, Gerdt C, Brokaw LP, Ros-Rocher N, Morris J, Davidson WS, Gerdt JP. Host lipids regulate multicellular behavior of a predator of a human pathogen. bioRxiv. 2024:2024.01.31.578218; doi: 10.1101/2024.01.31.578218.

9. Brunet T, King N. The Origin of Animal Multicellularity and Cell Differentiation. Dev Cell. 2017;43(2):124–40; doi: 10.1016/j.devcel.2017.09.016.

10. Ruiz-Trillo I, Kin K, Casacuberta E. The Origin of Metazoan Multicellularity: A Potential Microbial Black Swan Event. Annual Review of Microbiology. 2023;77(Volume 77, 2023):499–516; doi: 10.1146/annurev-micro-032421-120023.

11. Ereskovsky A, Borisenko IE, Bolshakov FV, Lavrov AI. Whole-Body Regeneration in Sponges: Diversity, Fine Mechanisms, and Future Prospects. Genes (Basel). 2021;12(4); doi: 10.3390/genes12040506.

12. Simunovic M, Metzger JJ, Etoc F, Yoney A, Ruzo A, Martyn I, Croft G, You DS, Brivanlou AH, Siggia ED. A 3D model of a human epiblast reveals BMP4-driven symmetry breaking. Nat Cell Biol. 2019;21(7):900–10; doi: 10.1038/s41556-019-0349-7.

13. McKeown SJ, Wallace AS, Anderson RB. Expression and function of cell adhesion molecules during neural crest migration. Dev Biol. 2013;373(2):244–57; doi: 10.1016/j.ydbio.2012.10.028.

14. O’Flaherty JT, Ward PA. Leukocyte aggregation induced by chemotactic factors. Inflammation. 1978;3(2):177–94; doi: 10.1007/BF00910738.

15. Woznica A, Gerdt JP, Hulett RE, Clardy J, King N. Mating in the Closest Living Relatives of Animals Is Induced by a Bacterial Chondroitinase. Cell. 2017;170(6):1175–83.e11; doi: 10.1016/j.cell.2017.08.005.

16. Ros-Rocher N, Reyes-Rivera J, Foroughijabbari Y, Combredet C, Larson BT, Coyle MC, Houtepen EAT, Vermeij MJA, King N, Brunet T. Mixed clonal-aggregative multicellularity entrained by extreme salinity fluctuations in a close relative of animals. bioRxiv. 2024:2024.03.25.586565; doi: 10.1101/2024.03.25.586565.

17. Sebé-Pedrós A, Peña Marcia I, Capella-Gutiérrez S, Antó M, Gabaldón T, Ruiz-Trillo I, Sabidó E. High-Throughput Proteomics Reveals the Unicellular Roots of Animal Phosphosignaling and Cell Differentiation. Developmental Cell. 2016;39(2):186–97; doi: 10.1016/j.devcel.2016.09.019.

18. Kontush A, Lhomme M, Chapman MJ. Unraveling the complexities of the HDL lipidome. J Lipid Res. 2013;54(11):2950–63; doi: 10.1194/jlr.R036095.

19. Schumann-Gillett A, O’Mara ML. The effects of oxidised phospholipids and cholesterol on the biophysical properties of POPC bilayers. Biochimica et Biophysica Acta (BBA) - Biomembranes. 2019;1861(1):210–9; doi: 10.1016/j.bbamem.2018.07.012.

20. Marsh D. Handbook of Lipid Bilayers. 2nd ed. Boca Raton: CRC Press; 2013.

21. Goldstein JL, Anderson RG, Brown MS. Receptor-mediated endocytosis and the cellular uptake of low density lipoprotein. Ciba Found Symp. 1982(92):77–95; doi: 10.1002/9780470720745.ch5.

22. Ritter P, Yousefi K, Ramirez J, Dykxhoorn DM, Mendez AJ, Shehadeh LA. LDL Cholesterol Uptake Assay Using Live Cell Imaging Analysis with Cell Health Monitoring. J Vis Exp. 2018 (141); doi: 10.3791/58564.

23. Hetrick B, Han MS, Helgeson LA, Nolen BJ. Small molecules CK-666 and CK-869 inhibit actin-related protein 2/3 complex by blocking an activating conformational change. Chem Biol. 2013;20(5):701–12; doi: 10.1016/j.chembiol.2013.03.019.

24. Kirchhausen T, Macia E, Pelish HE. Use of dynasore, the small molecule inhibitor of dynamin, in the regulation of endocytosis. Methods Enzymol. 2008;438:77–93; doi: 10.1016/s0076-6879(07)38006-3.

25. Hill TA, Gordon CP, McGeachie AB, Venn-Brown B, Odell LR, Chau N, Quan A, Mariana A, Sakoff JA, Chircop M, Robinson PJ, McCluskey A. Inhibition of dynamin mediated endocytosis by the dynoles--synthesis and functional activity of a family of indoles. J Med Chem. 2009;52(12):3762–73; doi: 10.1021/jm900036m.

26. Quan A, McGeachie AB, Keating DJ, van Dam EM, Rusak J, Chau N, Malladi CS, Chen C, McCluskey A, Cousin MA, Robinson PJ. Myristyl trimethyl ammonium bromide and octadecyl trimethyl ammonium bromide are surface-active small molecule dynamin inhibitors that block endocytosis mediated by dynamin I or dynamin II. Mol Pharmacol. 2007;72(6):1425–39; doi: 10.1124/mol.107.034207.

27. Vercauteren D, Vandenbroucke RE, Jones AT, Rejman J, Demeester J, De Smedt SC, Sanders NN, Braeckmans K. The use of inhibitors to study endocytic pathways of gene carriers: optimization and pitfalls. Mol Ther. 2010;18(3):561–9; doi: 10.1038/mt.2009.281.

28. Fauzyah Y, Ono C, Torii S, Anzai I, Suzuki R, Izumi T, Morioka Y, Maeda Y, Okamoto T, Fukuhara T, Matsuura Y. Ponesimod suppresses hepatitis B virus infection by inhibiting endosome maturation. Antiviral Res. 2021;186:104999; doi: 10.1016/j.antiviral.2020.104999.

29. Wu Y, Pons V, Goudet A, Panigai L, Fischer A, Herweg JA, Kali S, Davey RA, Laporte J, Bouclier C, Yousfi R, Aubenque C, Merer G, Gobbo E, Lopez R, Gillet C, Cojean S, Popoff MR, Clayette P, Le Grand R, Boulogne C, Tordo N, Lemichez E, Loiseau PM, Rudel T, Sauvaire D, Cintrat JC, Gillet D, Barbier J. ABMA, a small molecule that inhibits intracellular toxins and pathogens by interfering with late endosomal compartments. Sci Rep. 2017;7(1):15567; doi: 10.1038/s41598-017-15466-7.

30. Gillespie EJ, Ho CL, Balaji K, Clemens DL, Deng G, Wang YE, Elsaesser HJ, Tamilselvam B, Gargi A, Dixon SD, France B, Chamberlain BT, Blanke SR, Cheng G, de la Torre JC, Brooks DG, Jung ME, Colicelli J, Damoiseaux R, Bradley KA. Selective inhibitor of endosomal trafficking pathways exploited by multiple toxins and viruses. Proc Natl Acad Sci U S A. 2013;110(50):E4904–12; doi: 10.1073/pnas.1302334110.

31. Al-Bari MAA. Targeting endosomal acidification by chloroquine analogs as a promising strategy for the treatment of emerging viral diseases. Pharmacol Res Perspect. 2017;5(1):e00293; doi: 10.1002/prp2.293.

32. Misinzo G, Delputte PL, Nauwynck HJ. Inhibition of endosome-lysosome system acidification enhances porcine circovirus 2 infection of porcine epithelial cells. J Virol. 2008;82(3):1128–35; doi: 10.1128/jvi.01229-07.

33. Darch SE, West SA, Winzer K, Diggle SP. Density-dependent fitness benefits in quorum-sensing bacterial populations. Proc Natl Acad Sci U S A. 2012;109(21):8259–63; doi: 10.1073/pnas.1118131109.

34. Skotland T, Sagini K, Sandvig K, Llorente A. An emerging focus on lipids in extracellular vesicles. Advanced Drug Delivery Reviews. 2020;159:308–21; doi: 10.1016/j.addr.2020.03.002.

35. Koga Y, Kusaka I. Involvement of Autolysis of Cytoplasmic Membranes in the Process of Autolysis of Bacillus cereus. Microbiology. 1968;53(2):253–8; doi: 10.1099/00221287-53-2-253.

36. Esch BM, Fröhlich F. Chapter One - Mechanisms of Lipid Sorting in the Endosomal Pathway. In: Iglič A, Rappolt M, García-Sáez AJ, editors. Advances in Biomembranes and Lipid Self-Assembly: Academic Press; 2018. p. 1–39.

37. Lane-Donovan C, Philips GT, Herz J. More than cholesterol transporters: lipoprotein receptors in CNS function and neurodegeneration. Neuron. 2014;83(4):771–87; doi: 10.1016/j.neuron.2014.08.005.

38. May P, Herz J, Bock HH. Molecular mechanisms of lipoprotein receptor signalling. Cellular and Molecular Life Sciences CMLS. 2005;62(19):2325–38; doi: 10.1007/s00018-005-5231-z.

39. Mineo C. Lipoprotein receptor signalling in atherosclerosis. Cardiovasc Res. 2020;116(7):1254–74; doi: 10.1093/cvr/cvz338.

40. Schindelin J, Arganda-Carreras I, Frise E, Kaynig V, Longair M, Pietzsch T, Preibisch S, Rueden C, Saalfeld S, Schmid B, Tinevez J-Y, White DJ, Hartenstein V, Eliceiri K, Tomancak P, Cardona A. Fiji: an open-source platform for biological-image analysis. Nature Methods. 2012;9(7):676–82; doi: 10.1038/nmeth.2019.

41. Chernin E. Observations on hearts explanted in vitro from the snail Australorbis glabratus. J Parasitol. 1963;49:353–64

42. Developers F. ffmpeg tool. ffmpeg-20200213-6d37ca8-win64-static ed: Free Software Foundation, Inc.; 2007. p. 2020 Build by Kyle Schwarz.

## References

1. von Kleist L, Stahlschmidt W, Bulut H, Gromova K, Puchkov D, Robertson Mark J, MacGregor Kylie A, Tomilin N, Pechstein A, Chau N, Chircop M, Sakoff J, von Kries Jens P, Saenger W, Kräusslich H-G, Shupliakov O, Robinson Phillip J, McCluskey A, Haucke V. Role of the Clathrin Terminal Domain in Regulating Coated Pit Dynamics Revealed by Small Molecule Inhibition. Cell. 2011;146(3):471–84; doi: 10.1016/j.cell.2011.06.025.

2. Hill TA, Gordon CP, McGeachie AB, Venn-Brown B, Odell LR, Chau N, Quan A, Mariana A, Sakoff JA, Chircop M, Robinson PJ, McCluskey A. Inhibition of dynamin mediated endocytosis by the dynoles--synthesis and functional activity of a family of indoles. J Med Chem. 2009;52(12):3762–73; doi: 10.1021/jm900036m.

3. Kirchhausen T, Macia E, Pelish HE. Use of dynasore, the small molecule inhibitor of dynamin, in the regulation of endocytosis. Methods Enzymol. 2008;438:77–93; doi: 10.1016/s0076-6879(07)38006-3.

4. Quan A, McGeachie AB, Keating DJ, van Dam EM, Rusak J, Chau N, Malladi CS, Chen C, McCluskey A, Cousin MA, Robinson PJ. Myristyl trimethyl ammonium bromide and octadecyl trimethyl ammonium bromide are surface-active small molecule dynamin inhibitors that block endocytosis mediated by dynamin I or dynamin II. Mol Pharmacol. 2007;72(6):1425–39; doi: 10.1124/mol.107.034207.

5. Hetrick B, Han MS, Helgeson LA, Nolen BJ. Small molecules CK-666 and CK-869 inhibit actin-related protein 2/3 complex by blocking an activating conformational change. Chem Biol. 2013;20(5):701–12; doi: 10.1016/j.chembiol.2013.03.019.

6. Vercauteren D, Vandenbroucke RE, Jones AT, Rejman J, Demeester J, De Smedt SC, Sanders NN, Braeckmans K. The use of inhibitors to study endocytic pathways of gene carriers: optimization and pitfalls. Mol Ther. 2010;18(3):561–9; doi: 10.1038/mt.2009.281.

7. Fauzyah Y, Ono C, Torii S, Anzai I, Suzuki R, Izumi T, Morioka Y, Maeda Y, Okamoto T, Fukuhara T, Matsuura Y. Ponesimod suppresses hepatitis B virus infection by inhibiting endosome maturation. Antiviral Res. 2021;186:104999; doi: 10.1016/j.antiviral.2020.104999.

8. Wu Y, Pons V, Goudet A, Panigai L, Fischer A, Herweg JA, Kali S, Davey RA, Laporte J, Bouclier C, Yousfi R, Aubenque C, Merer G, Gobbo E, Lopez R, Gillet C, Cojean S, Popoff MR, Clayette P, Le Grand R, Boulogne C, Tordo N, Lemichez E, Loiseau PM, Rudel T, Sauvaire D, Cintrat JC, Gillet D, Barbier J. ABMA, a small molecule that inhibits intracellular toxins and pathogens by interfering with late endosomal compartments. Sci Rep. 2017;7(1):15567; doi: 10.1038/s41598-017-15466-7.

9. Gillespie EJ, Ho CL, Balaji K, Clemens DL, Deng G, Wang YE, Elsaesser HJ, Tamilselvam B, Gargi A, Dixon SD, France B, Chamberlain BT, Blanke SR, Cheng G, de la Torre JC, Brooks DG, Jung ME, Colicelli J, Damoiseaux R, Bradley KA. Selective inhibitor of endosomal trafficking pathways exploited by multiple toxins and viruses. Proc Natl Acad Sci U S A. 2013;110(50):E4904–12; doi: 10.1073/pnas.1302334110.

10. Misinzo G, Delputte PL, Nauwynck HJ. Inhibition of endosome-lysosome system acidification enhances porcine circovirus 2 infection of porcine epithelial cells. J Virol. 2008;82(3):1128–35; doi: 10.1128/jvi.01229-07.

